# A cautionary note on the use of haplotype callers in Phylogenomics

**DOI:** 10.1101/2020.06.10.145011

**Authors:** Pablo Duchen, Nicolas Salamin

## Abstract

Next-generation-sequencing haplotype callers are commonly used in studies to call variants from newly-sequenced species. However, due to the current availability of genomic resources, it is still common practice to use only one reference genome for a given genus, or even one reference for an entire clade of a higher taxon. The problem with traditional haplotype callers such as the one from GATK, is that they are optimized for variant calling at the population level, but not at the phylogenetic level. Thus, the consequences for downstream analyses can be substantial. Here, through simulations, we compare the performance between the haplotype callers of GATK and ATLAS, and present their differences at various phylogenetic scales. We show how the haplotype caller of GATK substantially underestimates the number of variants at the phylogenetic level, but not at the population level. We also quantified the level at which the accuracy of heterozygote calls declines with increasing distance to the reference genome. Such decrease is very sharp in GATK, while ATLAS maintains a high accuracy in variant calling, even at moderately-divergent species from the reference. We further suggest that efforts should be taken towards acquiring more reference genomes per species, before pursuing high-scale phylogenomic studies.

## 1 Introduction

Next-generation sequencing (NGS) grants access to a wealth of genomic data such as full genomes, transcriptomes, enriched target loci or RADseq. The increased availability of such data has revolutionized the way we study evolutionary processes at the population and species levels. Consequently, the growth in technologies to improve the sequencing of vast amounts of genomic regions was accompanied by the development of a diverse range of bioinformatic tools to analyse this new data (McCormack et al., 2013).

One of the main constraints of current NGS approaches is the reliance on a reference genome to call genetic variants. Many model organisms (such as humans or the fruit fly *Drosophila melanogaster*) have well-annotated reference genomes (dos Santos et al., 2015). For non-model organisms, however, access to a species-specific reference genome is still very often limited. Nielsen et al. (2011) stated that for an appropriate variant-calling accuracy, the amount of sequence identity between the reads and the reference (or the amount of tolerable mismatches between reads and reference) has to be optimized for every species individually. For example, if the same optimization used for humans is used for more variable organisms there will be a significant loss of sequencing depth, and thus, variability in many regions will be underestimated (Nielsen et al., 2011). The effect of the reference genome used for the analysis of newly-sequenced individuals will therefore be different depending on the evolutionary scale being studied. Consequently, variant calling at a population level (with reference genomes close to the target populations) might be more accurate than variant calling at a phylogenetic level (with reference genomes that are distant to the species analysed). So far, there are no studies that have looked at this specific question.

What is also clear is that currently, we are still far from having well-annotated reference genomes for all species. For instance, countless studies over the past few years had access to only one single reference genome, if any, despite analysing multiple species within entire genera. Such is the case for fish (e.g. Chakrabarty et al., 2017; Burress et al., 2018; Hulsey et al., 2017, 2018), mammals (e.g. Kumar et al., 2017; Lima et al., 2018; Moura et al., 2020), birds (e.g. Ottenburghs et al., 2016), reptiles (e.g. Bragg et al., 2016), amphibians (e.g. Portik et al., 2016), insects (e.g. Yan et al., 2020), and plants (e.g. Nobre et al., 2018; Kreuzer et al., 2019; Helmstetter et al., 2020; Olvera-Mendoza et al., 2020; Brandrud et al., 2020; Wang et al., 2020). The situation can sometimes be even more dramatic with a single reference genome available for entire tribes or subfamilies, encompassing multiple genera (e.g. Hulsey et al., 2017; Wang et al., 2017; Hulsey et al., 2018; Heckenhauer et al., 2019; Loiseau et al., 2019).

The current situation raises the question of how read mapping and variant calling are affected by the distance between the target species (or populations) and the reference genome. As stated by Nielsen et al. (2011), using optimized pipelines developed for one specific species often results in sub-optimal variant calling in another species. For instance, Schubert et al. (2014) acknowledge the fact that the widely-used *Genome Analysis Toolkit* GATK (DePristo et al., 2011) is not well suited for non-human organisms, since, among other things, it depends on external information such as known sets of variant sites (often unavailable for non-model organisms).

Some studies have already addressed the impact of single references on multi-species NGS analyses. For instance, mapping to a single reference genome within one genus versus using de-novo assembly can result in less variants being called by the reference-based method (Fitz-Gibbon et al., 2017). Second, using few references for multiple species confounds paralogy with homology after a full genome duplication, which in turn, affects phylogenetic reconstruction (Chakrabarty et al., 2017). Third, the further the phylogenetic distance from a reference genome the less efficient (as measured by the sequencing-depth coverage) the NGS pipelines become, as it is the case for target enrichment (Bragg et al., 2016) or transcriptome-based loci (Portik et al., 2016).

The efficiency in heterozygote calling at the phylogenetic level can also be undermined when the reference genome is too distant. Consequently, heterozygote positions are often discarded due to the uncertainty in heterozygote calls, which in turn affects downstream analyses, such as the inference of divergence times (Lischer et al., 2014). Such problems also pertain to ploidy levels above two, for which the development of newer methods for variant calling are necessary (Blischak et al., 2018). Finally, another important obstacle when calling variants in a newly-sequenced species is the lack of knowledge on species-specific variable sites, which are used to re-calibrate base quality scores (Schubert et al., 2014). Such knowledge is very helpful for judging the accuracy of variant calls, and for obtaining more accurate SNPs.

Overall, the degree of information loss when using one reference genome for multi-species NGS studies is currently unknown. Therefore, we propose here a simulation study to assess the efficiency of variant calling across various phylogenetic scales. We hypothesise a sharp drop in this efficiency as the evolutionary distance to the reference genome increases. The advantage of using simulations is that we know the exact position of variable sites, so that variant calling using traditional pipelines can be tested and the loss of information quantified. More specifically, we simulated diploid reads along different tree topologies and different evolutionary scales. For every tree and every scaling, we selected one reference genome and performed variant calling with the haplotype callers of GATK and ATLAS (Link et al., 2017). We focused on two aspects that affect downstream analyses substantially, namely: the number of called variants, and the accuracy of heterozygote calling.

## 2 Methods

Our pipeline followed these general steps: simulations of coalescent and phylogenetic trees, re-scaling of the trees to match various levels of divergence, simulation of genomic sequences along the trees, and simulation of diploid reads from each genomic sequence. From each tree, we selected one sequence as reference and performed standard genome assembly and variant calling with GATK 4.1 and ATLAS. Each of these steps is detailed below.

### 2.1 Simulation of trees and scaling

We performed two separate sets of tree simulations and scalings. As a first set, we simulated two contrasting topologies with arbitrary re-scalings of each topology (*Base cases* step). The motivation for the *base cases* analysis was to explore the performance of haplotype callers at varying phylogenetic scales while keeping the topology constant. For the second set of simulations (*Generalization* step), we simulated 10 random topologies and drew random re-scaling values from a uniform distribution. The motivation for this latter analysis was to check the robustness of our results with several randomly-generated topologies.

#### Base cases

As stated above, we used two types of trees (each with 20 tips) representing two contrasting topologies. The purpose of using these two topologies was to make an initial assessment of the potential effect that the topology and the position of the reference genome can have on haplotype calling. The first tree was a standard birth-death tree simulated with the R package *TreeSim* by Stadler (2011) (we called this “Tree A”, Fig. 1, left panel). The second tree was a highly-structured tree simulated with *ms* (Hudson, 2002) (from now on “Tree B”, Fig. 1, right panel). We scaled the trees to three arbitrary total depths of 0.13, 0.065, and 0.013. Although these values were arbitrarily chosen, they do correspond to realistic values observed in phylogenetic trees at the genus level. For instance, the first scaling corresponds to the actual divergence observed in the phylogenetic tree of clownfishes (Litsios et al., 2014). The second and third scaling values correspond to half and one-tenth the original value of the first scaling, representing smaller clades within a phylogenetic tree. We called these re-scaled values Large, Medium, and Small, respectively. Thus, the simulations in the *base cases* step included six types of trees: tree A Large, tree A Medium, tree A Small, tree B Large, tree B Medium, and tree B Small (Table 1, first column). Finally, to check the performance of haplotype callers on a level of divergence that can be considered as including several distinct populations represented by single individuals, we added one more scaling of trees A and B with a much smaller divergence (tree depth of 0.0013).

**Figure 1:**
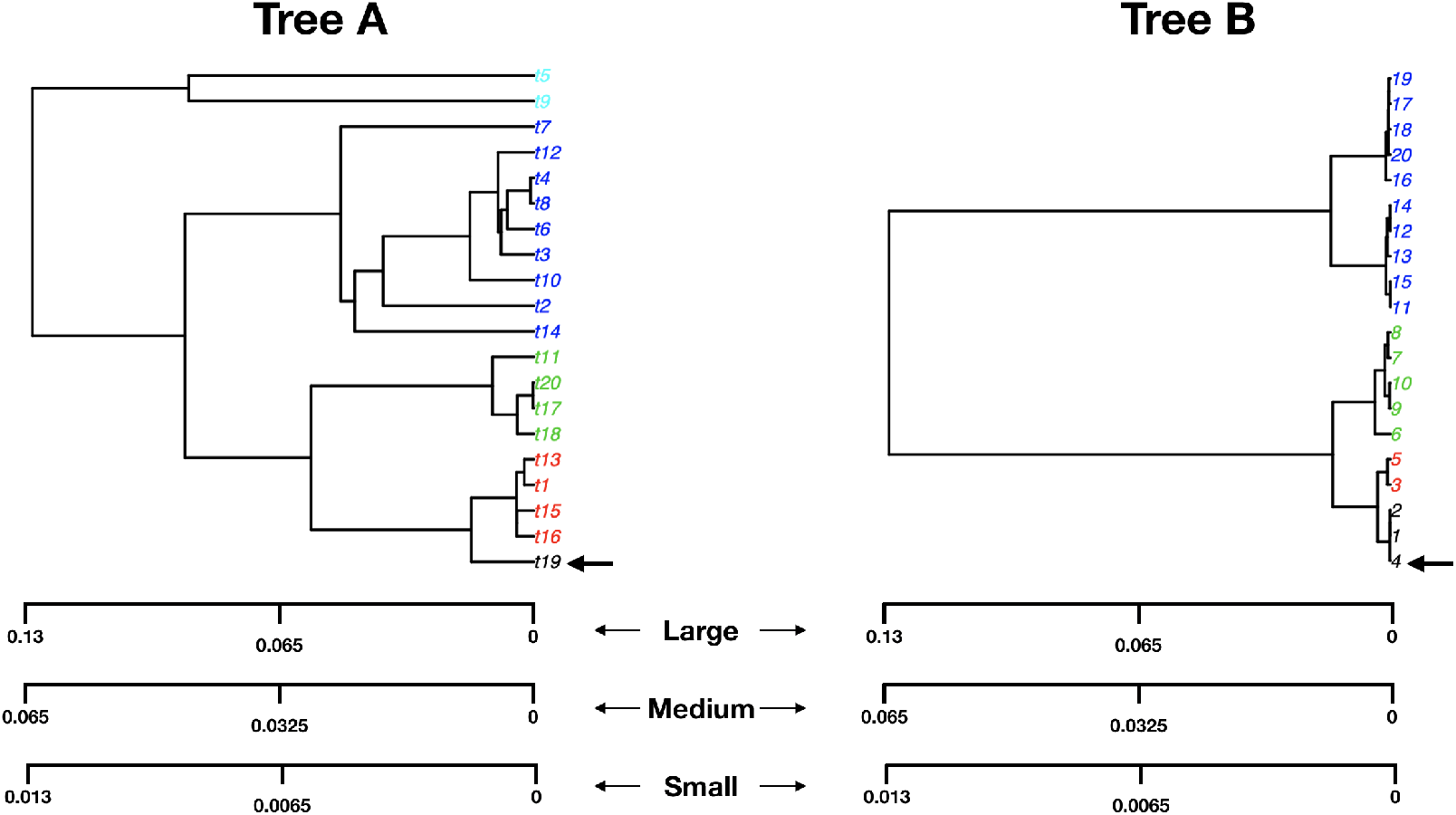
*Base-cases* example trees: a birth-death tree A (left), and a structured tree B (right). For each tree we generated three arbitrary re-scalings: large, medium, and small, each representing various types of divergence found in phylogenetic studies. The chosen reference genome is shown with an arrow. The colors of the tip labels are chosen to represent the increasing distance to the reference (these colors will be used again in Figures 3 and 4).

**Table 1:** Some stats from the sequence simulations, mapping of reads, and haplotype calling subroutines: Total number of SNPs for each alignment of each tree, mean (and standard deviation, sd) read-mapping quality (PHRED scores) for each tree (a per-species distribution of read-mapping qualities is shown in SI-Fig. B.1, and B.2), and total variant-calling computation time for all 20 species of each tree.

| Tree | Number of SNPs | Mean and (sd) PHRED | Computation time (GATK) | Computation time (ATLAS) |
| --- | --- | --- | --- | --- |
| A-large | 193884 | 57.04 (6.84) | ~ 6 hours | ~ 5 min. |
| A-medium | 97087 | 59.85 (1.47) | ~ 5 hours | ~ 4 min. |
| A-small | 19287 | 60.00 (0.00) | ~ 30 min. | ~ 4 min. |
| B-large | 143715 | 57.27 (6.83) | ~ 4 hours | ~ 3 min. |
| B-medium | 72113 | 59.74 (1.99) | ~ 3.5 hours | ~ 3 min. |
| B-small | 14256 | 60.00 (0.00) | ~ 30 min. | ~ 2 min. |
| A-population | 1978 | 60.00 (0.00) | ~ 10 min. | ~ 2 min. |
| B-population | 1429 | 60.00 (0.00) | ~ 10 min. | ~ 2 min. |

#### Generalization

A second series of simulations was performed by generating (using again the R package *TreeSim*) 10 random topologies of 20 species each, with a lineage birth rate drawn from the uniform distribution 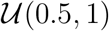 and an extinction rate drawn from 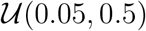. The trees were then re-scaled by drawing a scaling factor from a uniform distribution 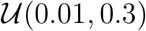. Such distribution spans divergence values both lower and higher than the values used in the *base cases* section.

### 2.2 Simulation of DNA sequences

For each of the trees generated in the previous section, we simulated 1 million bp-long sequences with the software *seq-gen* (Rambaut and Grass, 1997) under the HKY model (Hasegawa et al., 1985). In each tree, we repeated the simulations of sequences 20 times (one per tip) to account for the stochasticity that is inherent to models of sequence evolution. This will allow us to compare the different replicates and assess the variability introduced by this step of our simulation procedure. Concerning the expected percentage of invariant sites, we started with an arbitrary value of 80% for the large divergence scaling of trees A and B, which resulted in about 20% of variant sites in the final alignment. This percentage is in line with what can be observed in phylogenetic trees at the genus level (e.g. Duchen and Renner, 2010; Litsios et al., 2014). For the Medium and Small trees (with half and one-tenth of the divergence, respectively) we expect 10% and 2% of variant sites in the simulations of DNA sequences, assuming that mutation rates and all other simulation parameters are set to the same values (for a complete transcript of the pipeline and code please refer to the Supplementary Information). For the population-level tree, we expected to have around 0.2% of variant sites (provided that the original tree was 100 times more divergent and had 20% of variant sites). Finally, a similar approach was used for the *Generalization* step: the proportion of variable sites was taken between 2% and 20%, with the latter value assigned to the topology with the highest divergence.

### 2.3 Simulation of diploid reads

For each simulated sequence at the tips of every topology, we simulated 100,000 diploid reads (each read 100bp long) with the software *wg-sim* (Li, 2013). We kept the default parameters, which resulted in a total sequencing depth of 10x, a sequencing error rate of 0.001, and about 15% of indels introduced. We followed this approach because it allows us to know the exact position and state (heterozygote/homozygote) of all variants simulated, which is a key aspect that we want to test to understand the performance of haplotype callers.

### 2.4 Mapping of reads and generation of *bam* files

For every tree and scaling factor, we selected one sequence that was used as the reference for variant callling. In each simulated dataset, the reference sequence was indexed with *bwa index* (Li and Durbin, 2010), *samtools faidx* (Li et al., 2009), and *gatk CreateSequenceDictionary* (DePristo et al., 2011). The mapping of the simulated reads was done with *bwa* using the option *mem*. All *sam* files generated during this step were converted to *bam* format with *samtools view* and we discarded reads with mapping qualities below 30. We then sorted all *bam* files with the option *samtools sort*, added read groups with *gatk AddOrReplaceReadGroups*, and finally re-indexed the files with *samtools index*.

### 2.5 Base re-calibration and haplotype calling

Base re-calibration makes use of known variable positions. As recommended by the GATK Best Practices protocol (Van der Auwera et al., 2013), if these positions are unknown, one possible approach is to run the haplotype caller of GATK once (or a few times) and use the output to re-calibrate the quality scores. In our simulations, we know the positions of the variable sites but we did not use them for the re-calibration, since our goal was precisely to assess the performance of haplotype callers when there is no previous information on the positions of variable sites, which is the case in most newly-sequenced non-model organisms. We therefore ran the haplotype caller of GATK once to generate a table of inferred variable sites and the base re-calibration was then accomplished with the *BaseRecalibrator* and *ApplyBQSR* subroutines from GATK.

We used the *HaplotypeCaller* subroutine from GATK (DePristo et al., 2011), and the *MLE* subroutine of the software ATLAS (Link et al., 2017) to call haplotypes, using the exact same re-calibrated *bam* files described above as input in both cases. The main difference between ATLAS and GATK is that the former computes the genotype likelihoods of all ten possible genotypes at every given SNP, while the latter estimates the most likely minor allele before computing the three possible genotype likelihoods (see DePristo et al. (2011) and Link et al. (2017) for the description of both methods). Both haplotype callers generate VCF files (Danecek et al., 2011) as output. For each simulated SNP, we recorded whether the genotype returned by both callers matched the reference genome (i.e. “0/0” in the genotype field of the VCF file) or not (i.e. “1/1”), and whether the site was an heterozygote (i.e. “0/1”).

### 2.6 Performance of haplotype callers

#### 2.6.1 Total number of called variants

We measured the performance of the two haplotype callers GATK and ATLAS by first extracting the total number of called variants in every sequence of the simulated trees. This was done by counting in each VCF file the number of times a non “0/0” variant was called. We then compared this number to the true number of variants that is known from the simulations. This true number of variants can be estimated by counting the number of differences between each sequence and the corresponding reference, plus the number of simulated heterozygote positions, which is part of the output of the diploid-read simulator *wg-sim* (see section 2.3).

We focused on the number of variants for several reasons. First, the number of heterozygotes is an important measure of diversity, which is why it is expected for haplotype callers to correctly retrieve it, surpassing the potential effect of sequencing errors. Additionally, at the phylogenetic level, an accurate calling of homozygotes (that are different from the reference) can be an indicator of fixed substitutions that are, in turn, important to define the divergence between species and to measure the evolution of molecular markers.

#### 2.6.2 Accuracy in heterozygote variant calling

We extracted from the VCF files generated by GATK and ATLAS the positions of all heterozygote calls with the functions *extract.gt* and *is.het* from the R package *vcfR* (Knaus and Grünwald, 2017). We first tested if the total number of called heterozygotes corresponded to the true number of simulated heterozygotes. More precisely, we calculated the ratio between the number of called heterozygotes (by both GATK and ATLAS) and the true number of heterozygote positions obtained during the simulations. We called this ratio the *Number of Called/True heterozygotes*. We expected this ratio to be 1 for a perfect accuracy. A ratio above 1 would mean that there are more heterozygotes being called than the number simulated (representing false positives), and a ratio below 1 would mean that the number of heterozygote positions is underestimated by the haplotype caller. We also recorded if the positions of the called heterozygotes were correctly inferred by comparing them with the true positions simulated. This measure was estimated by counting the number of times a heterozygote position was correctly called by GATK and ATLAS and divided it by the total number of true heterozygote positions known from the simulated reads. We called this the *Accuracy of heterozygote variant calling*. We expected an accuracy of 1 for a perfect position match between called and true heterozygote positions.

## 3 Results

### 3.1 Simulation of sequences and reads

We initially simulated two topologies to illustrate the effect of the divergence between a reference genome and some new produced genomic data. These topologies were produced with a birth-death process (tree A) or a coalescent-based process (tree B), representing two contrasting tree shapes. We then re-scaled each topology to three different depths: Large (tree depth 0.13), Medium (tree depth 0.065), and Small (tree depth 0.013). We also added one more scaling of trees A and B to represent population-level divergences (tree depth 0.0013). We then simulated 10 additional topologies with birth and death rates taken from uniform distribution, and re-scaled each topology to depths between 0.01 and 0.3 (Fig. B.4, refer to section 2.1 for details). Simulation of reads yielded a total of 590 heterozygote positions per sequence (20 sequences were simulated per topology). The total number of segregating sites per alignment varied depending on the scale of the tree (Table 1).

### 3.2 Mapping of reads

The reads were mapped with different qualities depending on the distance to the corresponding reference (recall that we chose one reference per tree). For instance, reads belonging to the reference sequence or to tips adjacent to the reference had PHRED scores of 60. Mapping qualities then decreased with increasing distance to the reference (Fig. B.1, scale Large). Topologies of Medium and Small scales maintained an overall high read-mapping quality (Fig. B.1, scales Medium and Small). The same tendency was observed in the structured topology (tree B), with the only difference that we see a sharper difference in mapping quality when changing clades (Fig. B.2).

### 3.3 Haplotype calling

#### 3.3.1 Number of called variants

We counted the number of called variants, that is, all calls that are different from the reference (calls not of the form “0/0” in the genotype field of the output VCF file of each species). For the topology with the Small scale, we found that both GATK and ATLAS accurately recovered the number of variants (Fig. 2; third column). However, for Medium and Large scales GATK substantially underestimated the number of variants (Fig. 2; first and second columns). The caller ATLAS, on the other hand, did a perfect job for the Medium scales, but still underestimated the number of variants for the Large scale (particularly the tips that are far from the reference, Fig. 2, red lines), although this underestimation was not as sharp as the one seen for GATK. We observed a similar tendency for tree B (Fig. 2, second row). A similar pattern was also observed when we simulated random topologies with random scalings: when the divergence from the reference is low then both GATK and ATLAS recover the true number of variants, while, as soon as this divergence increased, GATK underestimated the number of variants, while ATLAS was able to outperform GATK (Fig. B.5).

**Figure 2:**
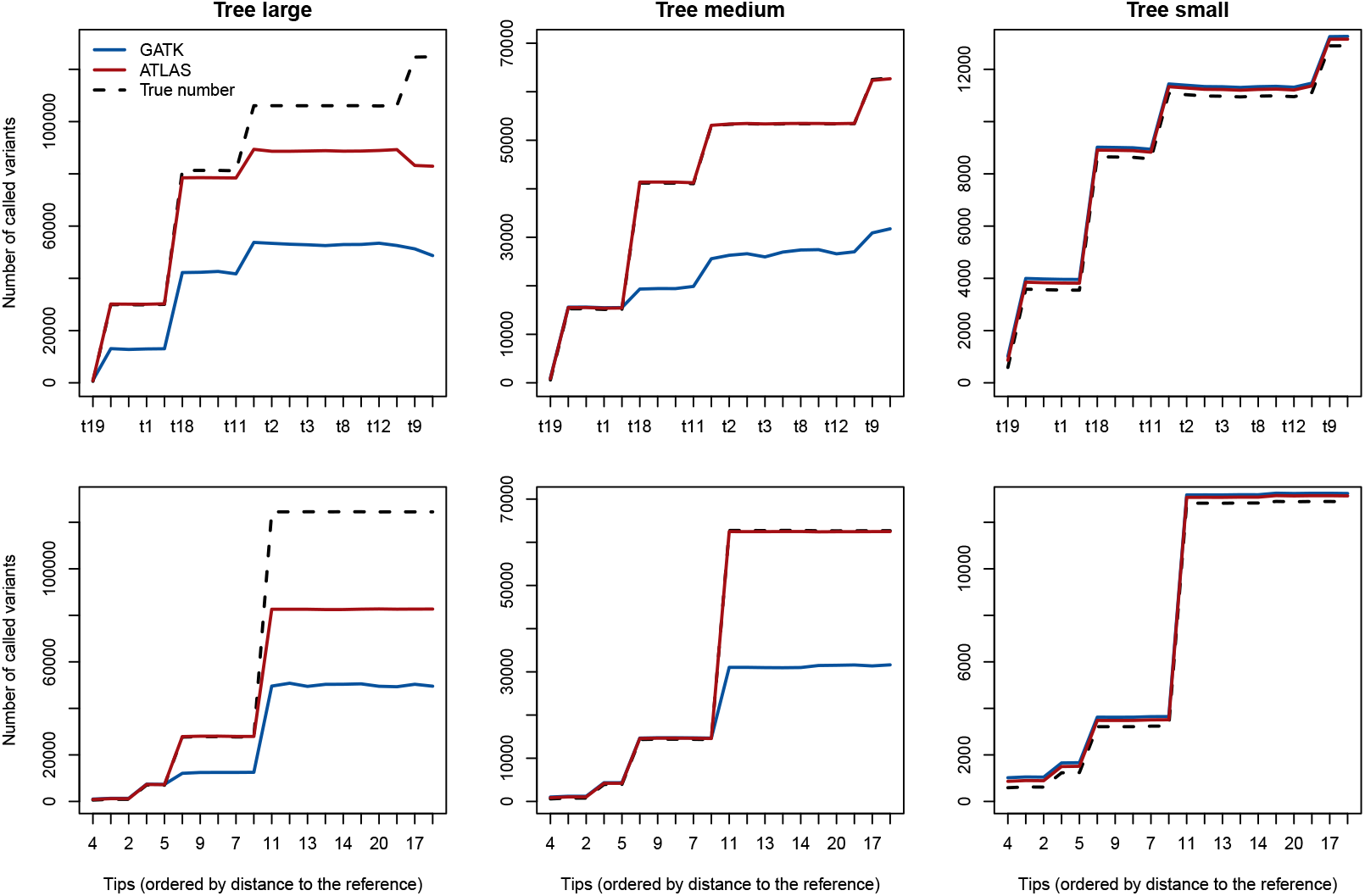
Number of called variants for the tips of tree A (first row), and tree B (second row). The tips on the x axis are ordered according to their distance to the reference (the reference being always at the extreme left).

#### 3.3.2 Calling of heterozygotes

After running the *HaplotypeCaller* and *MLE* subroutines of GATK and ATLAS respectively, we found little difference in their performance for the simulations at the population level. In this case, they both called the heterozygote variants accurately, although there was a 20% overestimation of the heterozygote calls by GATK (Fig. SI-B.3). However, we found the opposite result when dealing with the simulations at the phylogenetic-level: the accuracy in heterozygote calling reduced with increasing distance to the corresponding reference genome, and this reduction was particularly strong for GATK when compared to ATLAS. This pattern was observed in both types of topologies (Fig. 3 and 4, second column), as well as in the extra 10 simulated phylogenies under various scalings (Fig. B.6).

**Figure 3:**
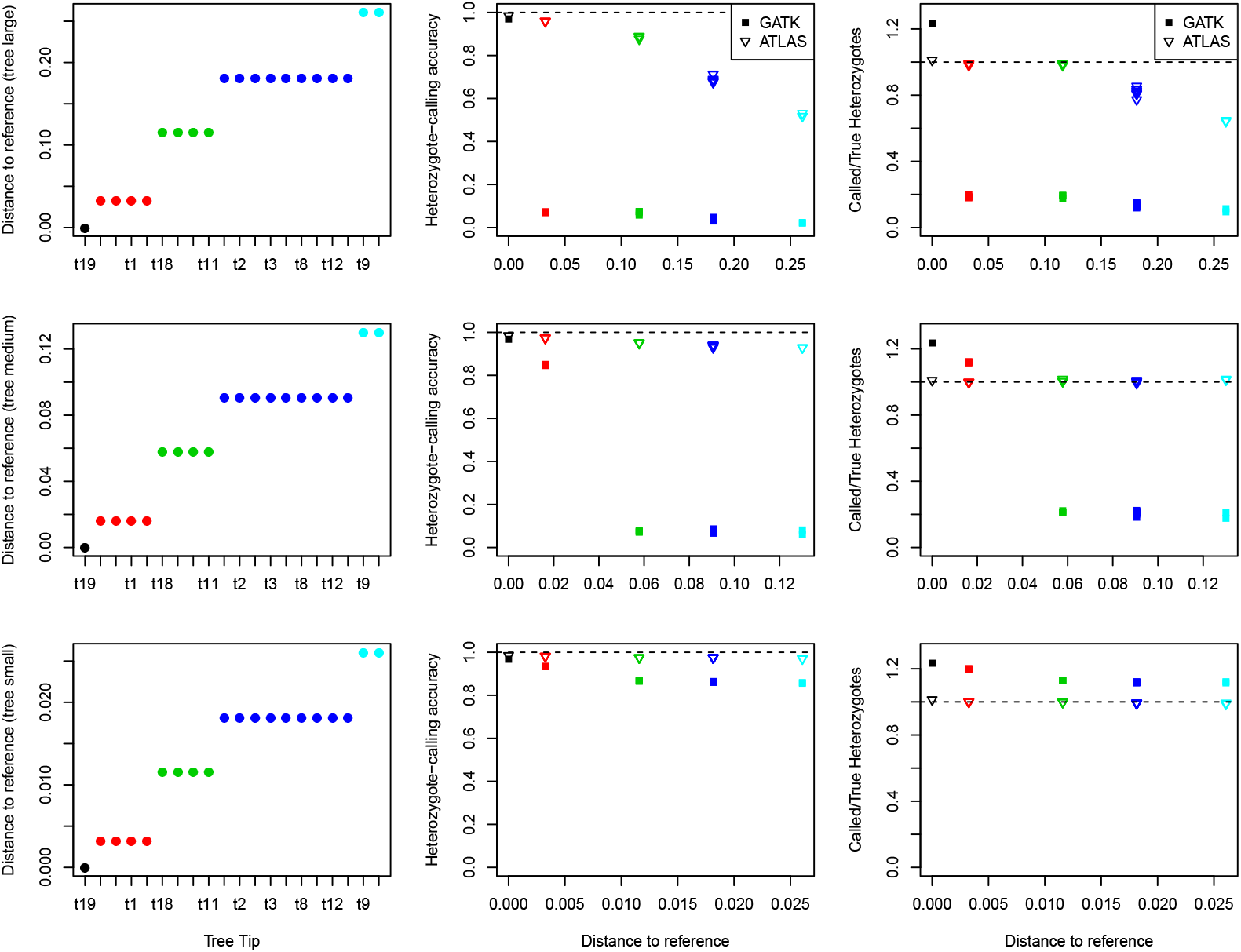
Accuracy of heterozygote calling for tree A at three different phylogenetic scales: large (first row), medium (middle row), and small (last row). The phylogenetic distance from each tip of the phylogeny to the reference is plotted against each tree tip (first column), against the heterozygote-calling accuracy (second column), and against the called versus true heterozygotes (third column). The colors of each point or symbol correspond to the tip colors shown in Fig. 1.

**Figure 4:**
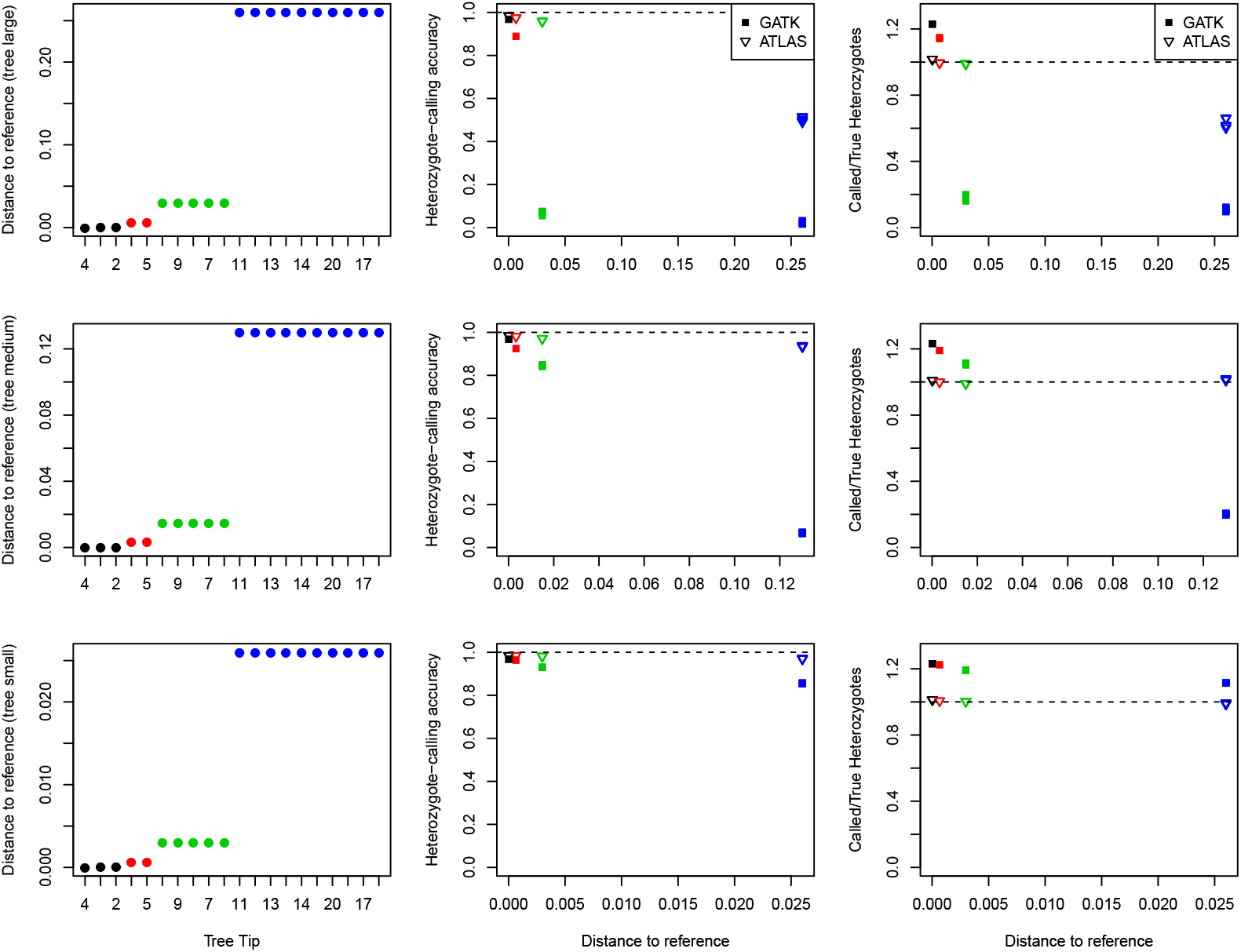
Accuracy of heterozygote calling for tree B at three different phylogenetic scales: large (first row), medium (middle row), and small (last row). The phylogenetic distance from each tip of the phylogeny to the reference is plotted against each tree tip (first column), against the heterozygote-calling accuracy (second column), and against the called versus true heterozygotes (third column). The colors of each point or symbol correspond to the tip colors shown in Fig. 1.

We also found that GATK tended to overestimate the number of heterozygotes when the genomes were close to the reference, but this tendency decreased dramatically with increasing phylogenetic distance to the reference. In other words, at a phylogenetic scale, GATK will substantially underestimate the true number of heterozygote positions along the genome. The haplotype caller of ATLAS, on the other hand, maintained the ratio of called vs true heterozygote variants close to 1 most of the time (Fig. 3 and 4, third column). A similar pattern was observed with the extra 10 simulated phylogenies under various scalings (Fig. B.7).

## 4 Discussion

In this study, we tested the performance of state-of-the-art haplotype callers on simulated datasets aimed at representing evolutionary divergences, either within or above the species level. Haplotype calling applied on reads simulated at a population level reached a good accuracy overall, although GATK overestimated the number of heterozygotes by as much as 20% (Fig. SI-B.3, last panel). Such overestimation of heterozygotes by GATK has also been reported in previous studies (Hwang et al., 2015; Link et al., 2017). In contrast, ATLAS was able to call heterozygotes with a 100% accuracy in all cases at the population level (Fig. SI-B.3, last panel). When the divergence between the simulated sequences increased to a level representative of phylogenomic studies, we found the performance of GATK sub-optimal. For instance, we observed a large underestimation in the total number of called variants (Fig. 2), and low accuracy when calling heterozygote positions (Fig. 3 and 4). While this issue is particularly prevalent in GATK, the caller ATLAS is less affected. The main difference between the two callers lies in their assumptions when estimating the possible genotype likelihoods necessary to call variants. ATLAS considers all ten possible genotype likelihoods, which might be better suited when divergent sequences are considered. GATK, on the other hand, infers one minor allele to be used in the genotype likelihood calculations. As expected, the under-performance of haplotype calling is stronger when the phylogenetic distance between the species-specific reads and the reference genome (against which these reads were mapped) becomes larger (see also Bragg et al., 2016; Portik et al., 2016; Fitz-Gibbon et al., 2017).

Depending on the research question, underestimating the number of variants can have different consequences, but failing to accurately call heterozy-gote positions can have more detrimental effects (Lischer et al., 2014), which is why we focused on this particular parameter in the present study. This problem can also be exacerbated for higher ploidy levels, for which newer methods for variant calling under such circumstances have been suggested (Blischak et al., 2018). Additionally, many studies have acknowledged the problems of lacking species-specific references, and they opted for the solution to build pseudo-references or synthetic references for every species instead (e.g. Bateman et al., 2018; Grummer et al., 2018; Skipwith et al., 2019). However, such studies focused more on target/sequence capture that are usually designed for interspecific studies and where the call of heterozygotes are of less interest. We focused instead on studies that make use of entire genomes for phylogenomic or population genomic analyses across multiple species, with well-annotated reference genomes.

The tree topology also seems to play a role in determining the accuracy of haplotype calling. Here, we started with two contrasting topologies: a “typical” birth-death topology and an extremely structured, coalescent-like topology (Fig. 1). In both cases, and at all scalings tested, the decrease of accuracy with increasing distance to the reference is not linear, but jumps as it goes through the different clades of the tree (Fig. 4, second and third columns). In other words, the further the most recent common ancestor between the reference and a given species is, the sharper the decrease in haplotype-calling accuracy that we found.

There are several reasons that might explain the low performance of haplotype calling when the divergence to the reference genome increases. First, the read mapping quality reduces with increasing distance to the reference because of the lower similarity between the reads and the reference sequences (Fig. SI-B.1 and SI-B.2). Second, the algorithms of the haplotype callers tested require a priori information about known variable sites to re-calibrate base quality scores (DePristo et al., 2011). However, this information is often unavailable for studies involving several species (Schubert et al., 2014). If this is the case, then we can expect that the species close to the reference might benefit from a better re-calibration than the species far from the reference.

In this study, we based our first scaling on the divergence that can be observed at the genus level for the clownfish phylogeny (Litsios et al., 2014). We are aware that, for other organisms, divergence at the genus level will be different, which is why we tested several additional scalings (Fig. 1 and Fig. B.4). We did not need to test larger divergence values because the performance of GATK on the trees simulated here already showed a rapid decrease with increasing distance to the reference. Given this trend, we expect this under-performance to be even greater for larger phylogenetic trees with greater divergence values among its species.

Nevertheless, we would like to make clear that current NGS software (such as GATK) does provide very good tools for NGS processing. It is only the haplotype-calling subroutine of GATK that needs to be used carefully, and applied only to population-level studies with a priori knowledge of variable sites, or use the ATLAS haplotype caller instead. We conclude that, for multi-species NGS studies, primary efforts should be taken towards building more reference genomes, rather than sequencing more species without a reference.

## Supplementary Information

### A General pipeline and code

#### A.1 Simulation and re-scaling of trees

The birth-death tree A was simulated with the R package *TreeSim* (Stadler, 2011), assuming both the birth and death rates are equal to 1:

~~~
library(TreeSim)
tree_A <-sim.bd.taxa(20,1,1,1,complete=FALSE)
write.tree(tree_A,file=“tree_A.tre”)
~~~

The *generalized* topologies were also simulated with *TreeSim* but with birth rates taken from the prior 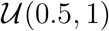, and death rates from the prior 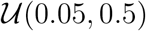.

The structured tree B was simulated with *ms* (Hudson, 2002):

~~~
./ms 20 1 -t 5 -T -I 8 5 0 5 0 5 0 5 0 \\
-ej 0.001 8 7 -ej 0.001 6 5 -ej 5 7 5 -ej 0.001 4 3 \\
-ej 0.001 2 1 -ej 5 3 1 -ej 50 5 1 \\
| grep ‘(’ > tree_B.tre)
~~~

Briefly, this command line simulates a tree with 20 tips divided in 4 clades, each composed of 5 tips. Commands “-ej” join clades at a given time in the past (e.g. -ej 0.001 8 7 joins clade 8 to clade 7 at 0.001 time units in the past).

Both trees A and B were re-scaled to tree depths of about 0.13, 0.065, and 0.013 with the function *rescale* of the R package *geiger* (Pennell et al., 2014):

~~~
library(geiger)
rescaled_tree <-rescale(tree,model=“depth”,new_depth)
~~~

### A.2 Simulation of sequences and diploid reads

Fasta sequences with a length of 1000000 bp were simulated with *seq-gen* (Rambaut and Grass, 1997) for each tree. For trees with the larger scale (depth of 0.13) we kept 80% of invariant sites, meaning that 20% of were SNPs (-i 0.8 as in the example below):

~~~
./seq-gen -m HKY -i 0.8 -l 1000000 -of -z 123456 tree_A.tre \\
> seqs_A.fa
~~~

For trees half this size or 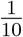 its size (depths of 0.065 and 0.013, respectively) we set the proportion of invariant sites accordingly (-i 0.9 and -i 0.98). For each sequence (20 sequences per tree) we simulated diploid reads with the program *wgsim* (Li, 2013) as follows:

~~~
./wgsim -N100000 -1100 -d0 -S11 -e0 -r0.001 \\
seqs_tip_1.fa reads_tip_1.fq /dev/null > output_wgsim_reads_tip_1.log
~~~

This command, plus its default settings, generates 100000 reads each with a length of 100bp, yielding an average sequencing depth of 10x, an error rate of 0.001, and about 15% of the total genome represented by indels.

### A.3 Mapping of simulated reads

From each tree we selected one tip sequence as the reference. Before mapping we indexed all references with *bwa* (Li and Durbin, 2010), *samtools* (Li et al., 2009), and GATK (DePristo et al., 2011) as follows:

~~~
./bwa index reference.fa
./samtools faidx reference.fa
./gatk CreateSequenceDictionary -R reference.fa
~~~

We then mapped all reads with the function *mem bwa*, and generated one SAM file per reads file, and then converted it to BAM format for downstream analyses:

~~~
./bwa mem reference.fa reads_tip_1.fq > reads_tip_1.sam
./samtools view -q 30 -S -b reads_tip_1.sam > reads_tip_1.bam
~~~

Then, we sorted the BAM files, added read groups with GATK, and indexed the output:

~~~
./samtools sort -o reads_tip_1_sorted.bam reads_tip_1.bam
./gatk AddOrReplaceReadGroups -I reads_tip_1_sorted.bam \\
-O reads_tip_1_sorted_RG.bam -RGID 4 -RGLB lib1 \\
-RGPL ILLUMINA -RGPU unit1 -RGSM 20
./samtools index reads_tip_1_sorted_RG.bam
~~~

Here, the suffix “RG” stands for “read group”.

### A.4 Base quality score recalibration

Before calling haplotypes, we generated a list of known sites for re-calibration with one round of the *HaplotypeCaller* routing of GATK:

~~~
./gatk HaplotypeCaller -I reads_tip_1_sorted_RG.bam -R reference.fa \\
-O known_sites.vcf
~~~

(Note: we performed this with only one sequence per tree, the sequence that was chosen as the reference).

Then, for every sequence in every tree we re-calibrated the quality scores with GATK as follows:

~~~
./gatk BaseRecalibrator -I reads_tip_1_sorted_RG.bam -R reference.fa \\
--known-sites known_sites.vcf -O reads_tip_1_sorted_RG.bam.table
./gatk ApplyBQSR -R reference.fa -I reads_tip_1_sorted_RG.bam \\
--bqsr-recal-file reads_tip_1_sorted_RG.bam.table \\
-O reads_tip_1_sorted_RG_RC.bam
~~~

Here, the suffix “RC” stands for “re-calibrated”.

### A.5 Haplotype calling

We called haplotypes with GATK and ATLAS (Link et al., 2017). The input for both programs was exactly the same, namely, the recalibrated bam files generated in the last step. For each read in each tree we used:

~~~
./gatk HaplotypeCaller -I reads_tip_1_sorted_RG_RC.bam -R reference.fa \\
-O reads_tip_1_sorted_RG_RC.vcf
~~~

for GATK, and for ATLAS we used:

~~~
./atlas task=call method=MLE bam=reads_tip_1_sorted_RG_RC.bam \\
fasta=reference.fa infoFields=DP formatFields=GT,AD,AB,AI,DP,GQ,PL
~~~

This last call with ATLAS generates one VCF file that is saved in the same folder as the input BAM file. In our particular run, we had to copy and rename all “*.bam.bai” files to “*.bai” to make sure they are read by ATLAS.

## B Supplementary Figures

**Figure B.1:**
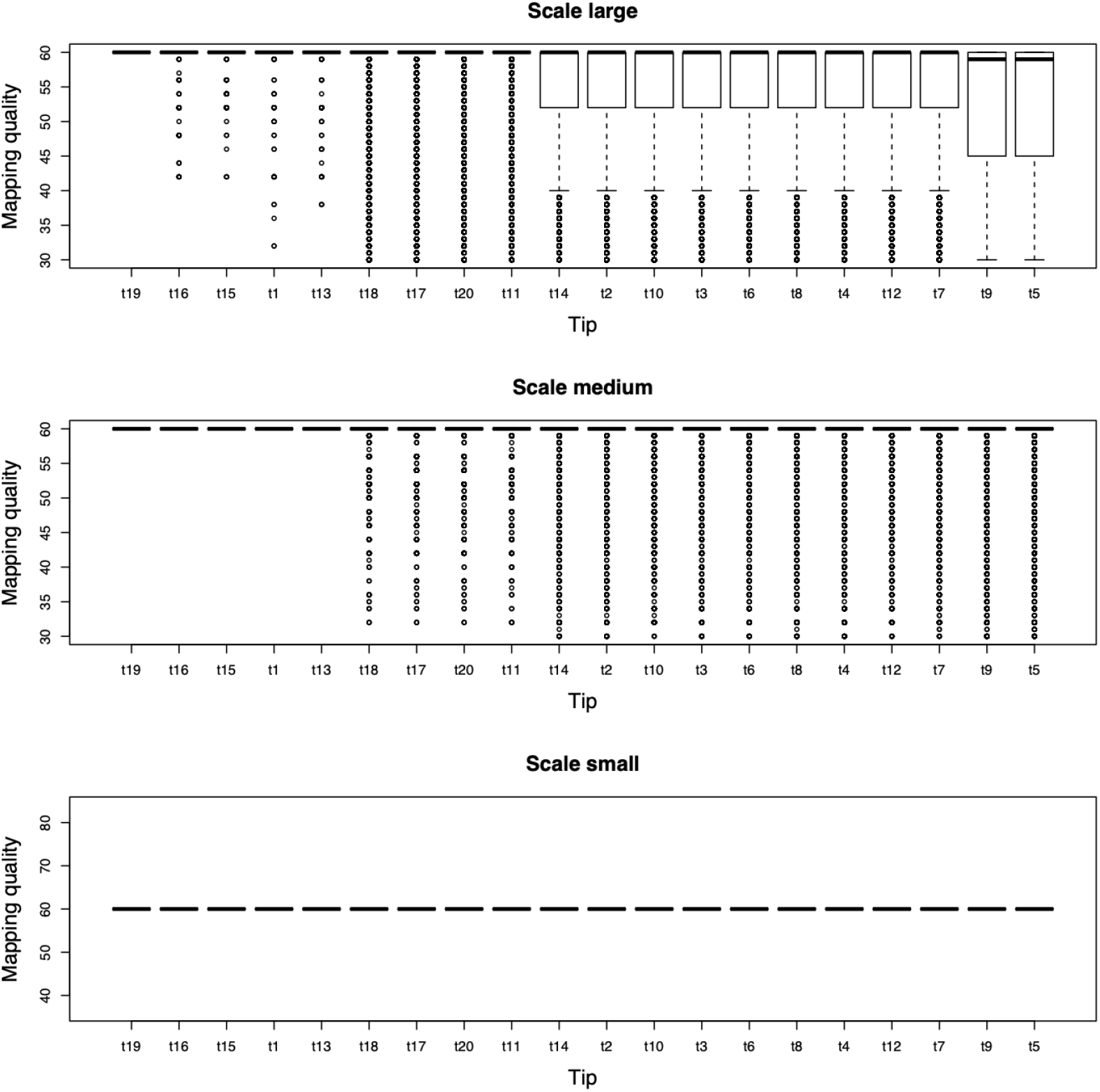
Mapping quality (PHRED score) reported in the SAM files generated by the *bwa* run for all tips of tree A, and for each scaling: large, medium, and small. Tips are ordered by increasing phylogenetic distance to the corresponding reference (the reference being at the extreme left).

**Figure B.2:**
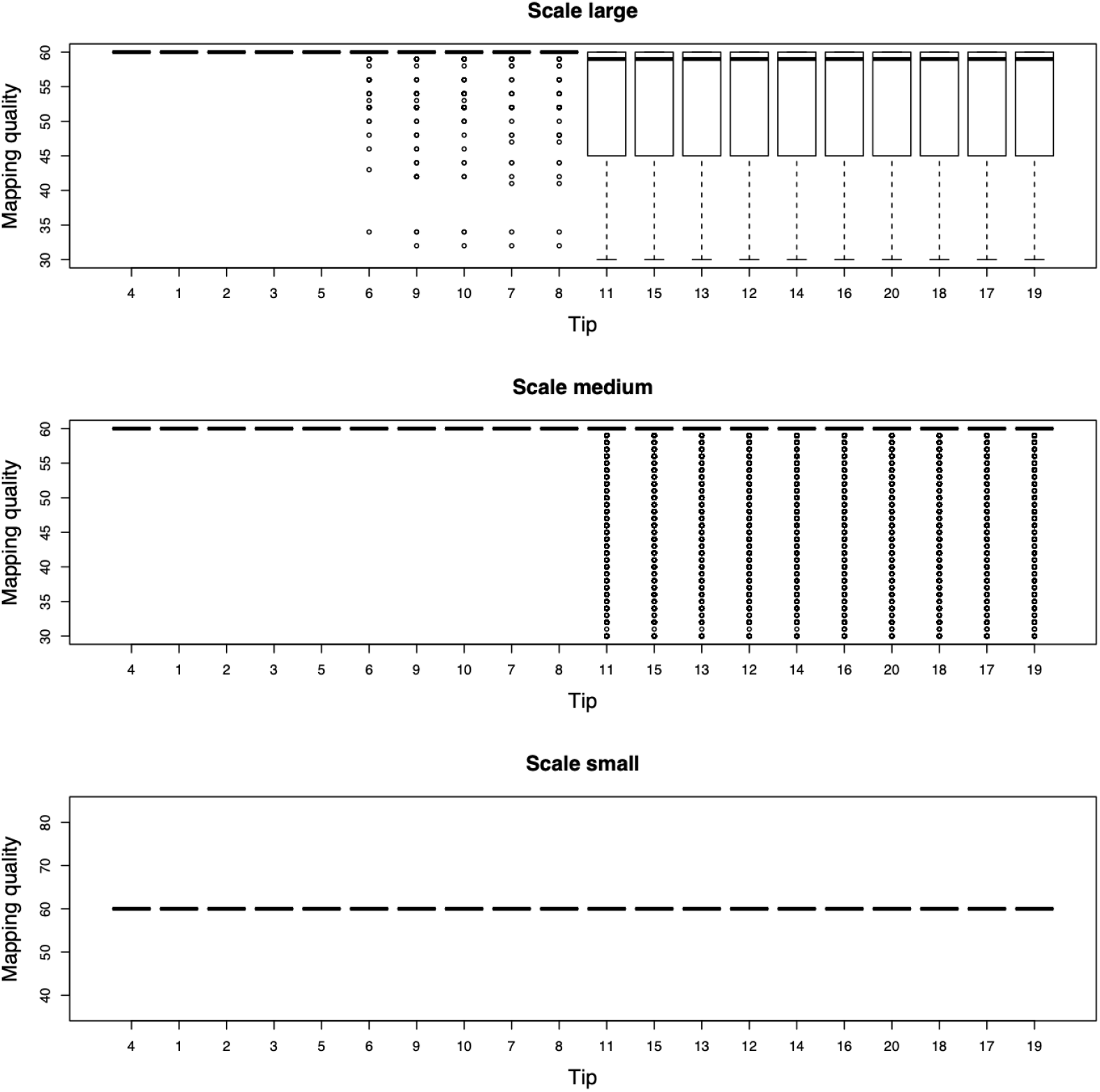
Mapping quality (PHRED score) reported in the SAM files generated by the *bwa* run for all tips of tree B, and for each scaling: large, medium, and small. Tips are ordered by increasing phylogenetic distance to the corresponding reference (the reference being at the extreme left).

**Figure B.3:**
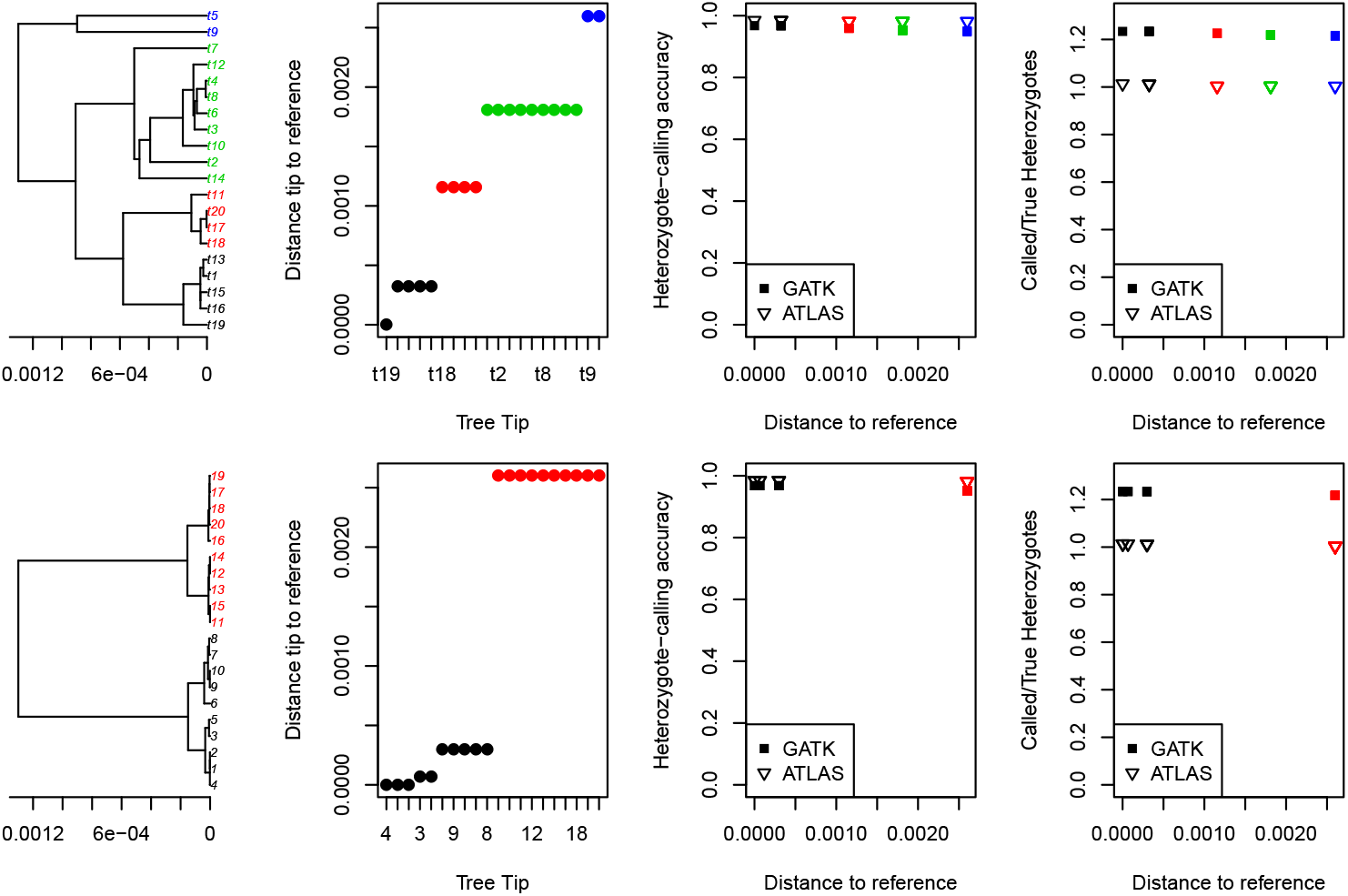
Accuracy of heterozygote-calling for two trees at a population-level scale (low divergence). Same as in Fig. 3 and 4, the phylogenetic distance from each tip of the phylogeny to the reference is plotted against each tree tip (second column), against the heterozygote-calling accuracy (third column), and against the called versus true heterozygotes (fourth column). The colors of each point or symbol correspond to the ones in the trees of the first column.

**Figure B.4:**
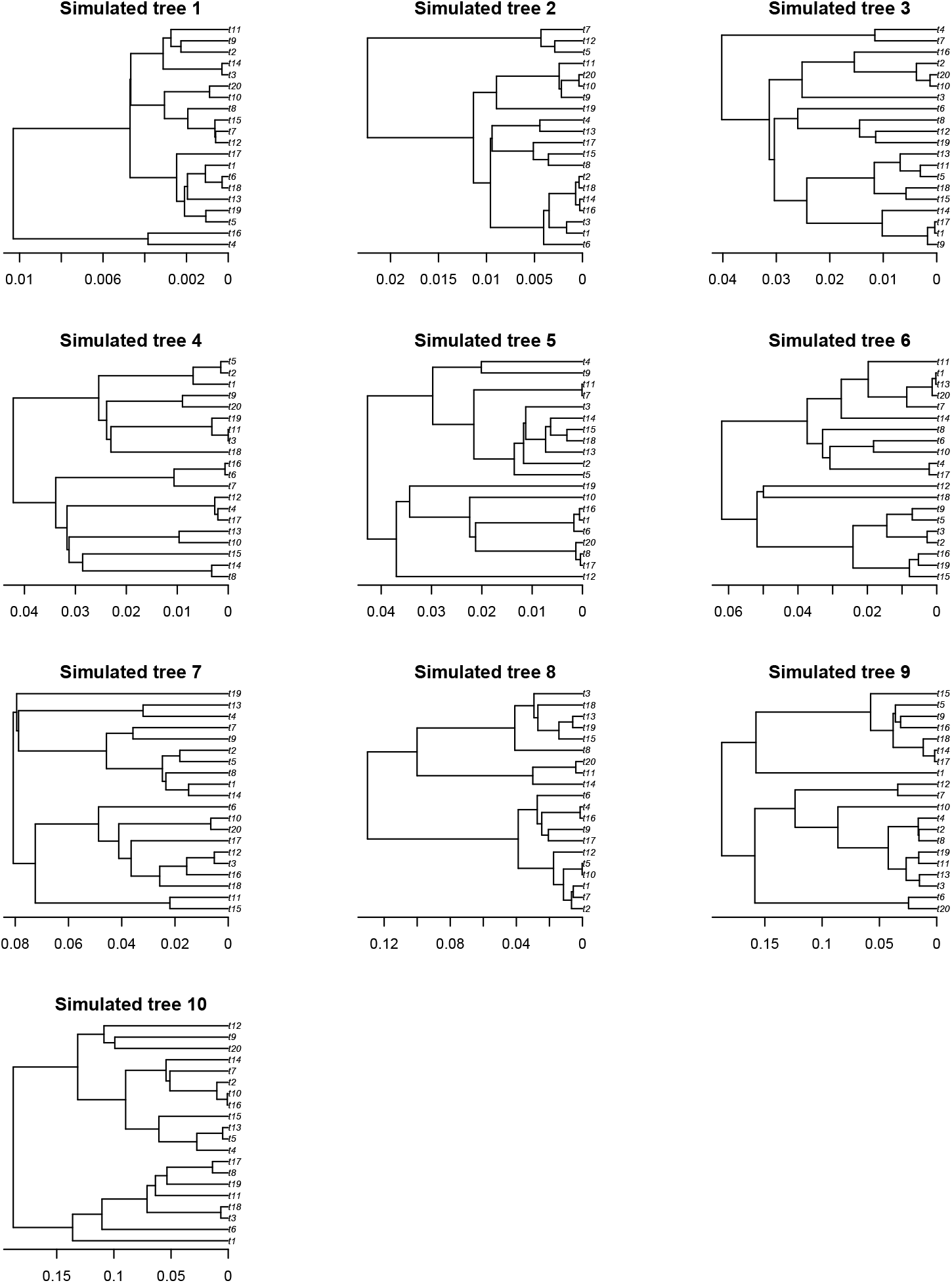
Simulated topologies and scalings used in the *generalization* step of our analysis (see section 2.1).Topologies were ordered by increasing divergence.

**Figure B.5:**
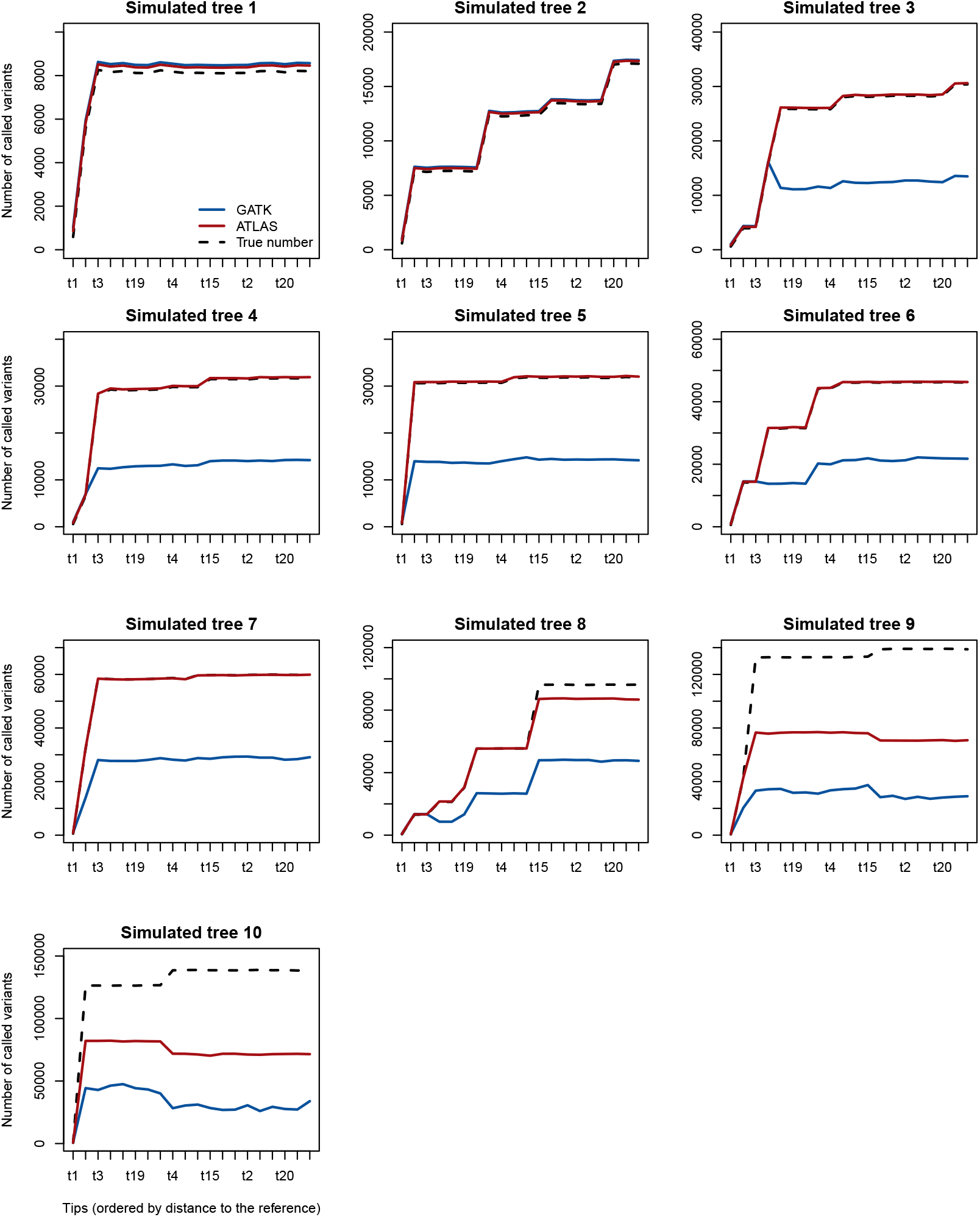
Number of called variants for the tips of the trees in Fig. B.4. The tips on the x axis are ordered according to their distance to the reference (the reference being always at the extreme left).

**Figure B.6:**
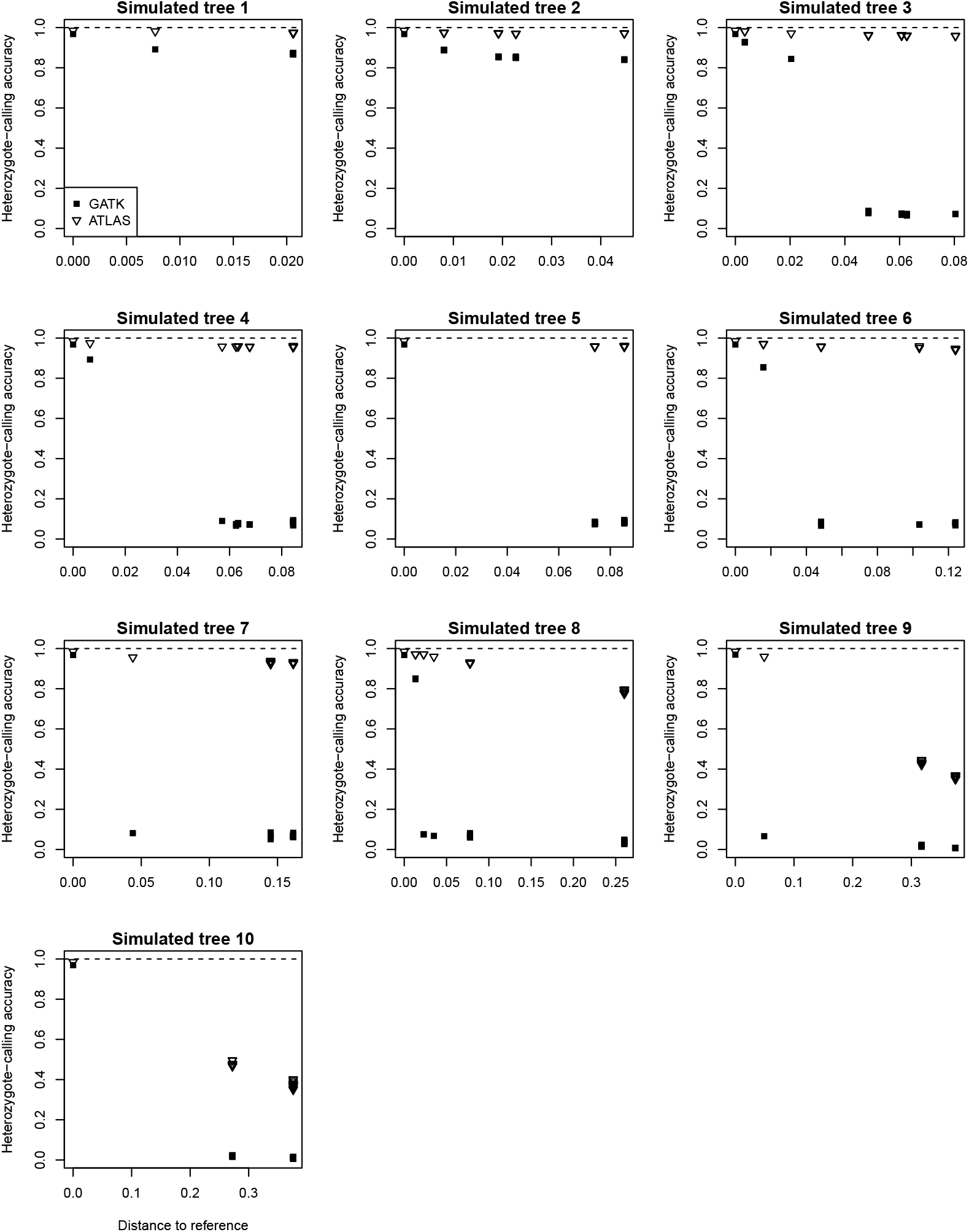
Accuracy of heterozygote calling for the trees in Fig. B.4. The phylogenetic distance is plotted against the heterozygote-calling accuracy.

**Figure B.7:**
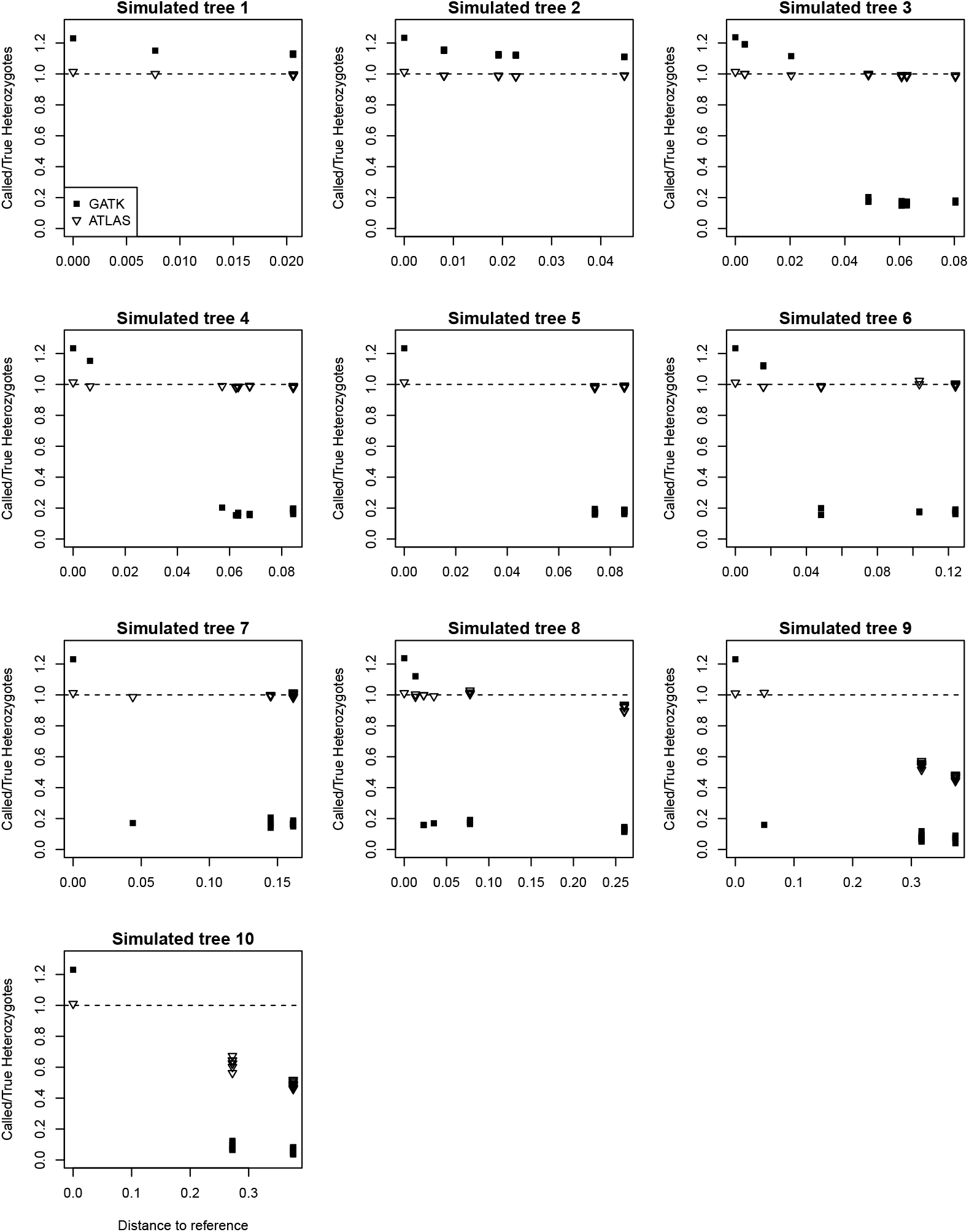
Accuracy of heterozygote calling for the trees in Fig. B.4. The phylogenetic distance is plotted against the called versus true heterozygotes.

## Notes

### Competing Interest Statement

The authors have declared no competing interest.

